# CausalKnowledgeTrace: A Novel Computational Framework for Automated Literature-Based Causal Graph Construction and Evidence-Based Variable Selection in Biomedical Research

**DOI:** 10.64898/2026.05.07.723601

**Authors:** Rajesh Upadhayaya, Manjil Pradhan, Vincent Metzger, Scott Alexander Malec

## Abstract

**Background:** Variable selection for causal inference from observational biomedical data is challenging, as overlooking confounders or conditioning on colliders leads to biased estimates. While vast causal knowledge exists in biomedical literature, manually extracting this information for principled variable selection is impractical at scale.

**Methods:** We developed CausalKnowledgeTrace, a Python-based computational framework with Django web interface that systematically leverages structured causal knowledge from the Semantic MEDLINE Database (SemMedDB) to inform variable selection in causal studies. The system implements a six-stage analysis pipeline using NetworkX for graph operations, including graph parsing, basic analysis, comprehensive cycle detection, systematic generic node removal, post-removal analysis, and formal causal inference with bias detection.

**Results:** Analysis of the hypertension-Alzheimer’s relationship across three degree neighborhoods (1-3) demonstrated systematic scaling of causal complexity: 361-866 variables, 429-1,442 relationships, with graph densities of 0.0033-0.0019. The analysis revealed complex cyclic structures with 54-606 baseline cycles across degree levels. Processing times ranged from 0.3-1.0 seconds for all three degrees, demonstrating computational efficiency for complex biomedical networks. Key confounders identified across all degrees included inflammation, diabetes, insulin resistance, obesity, and ischemia. In the third degree of graph, the pipeline structurally identified 39 confounders, 11 mediators, and 3 colliders from the causal graph. Among the key identified confounders and mediators—including obesity, oxidative stress, ischemia, and vascular diseases—all were found to have strong supporting evidence in established epidemiological and pathophysiological literature.

**Conclusions:** CausalKnowledgeTrace provides a scalable, evidence-based approach to causal graph construction that systematically identifies confounders and bias structures often missed by conventional approaches. The Python-Django architecture enables both standalone analysis and integration into larger computational workflows, representing a significant advance in computational support for causal inference in biomedical research.

**Statement of Significance:** *Problem or Issue:* Selecting proper confounders and variables for causal inference from observational biomedical datasets is challenging and often biased by limited expertise or manual review.

*What is Already Known:* Existing approaches rely on domain experts, statistical variable screening, or manual construction of causal graphs, but these often overlook literature-documented confounders and complex biases.

*What this Paper Adds:* This paper introduces an automated, literature-based framework for synthesizing and validating causal graphs, identifying critical variables and complex bias structures, such as M-bias and butterfly bias, with full evidentiary traceability.

*Who would benefit from the new knowledge in this paper?:* Epidemiologists, biomedical researchers, informaticians, and clinical investigators seeking reliable and transparent causal modeling for observational studies.

## 1. Introduction

Causal inference from observational data has become increasingly critical in biomedical research as the scale and complexity of available datasets continue to expand.^5^ Electronic health records, genomic databases, and large-scale epidemiological studies offer unprecedented opportunities to understand disease mechanisms, evaluate therapeutic interventions, and inform clinical decision-making.^4^ However, the validity of causal conclusions drawn from these observational data sources fundamentally depends on appropriate control for confounding variables—factors that influence both the exposure and the outcome of interest.^12^

The challenge of identifying appropriate confounding variables has intensified as biomedical research has evolved from small, focused studies to large-scale analyses involving hundreds of potential covariates. Traditional approaches to variable selection rely heavily on domain expertise, statistical screening methods, or manual literature review, each carrying significant limitations.^16^ Domain experts may overlook confounders documented in adjacent fields or emerging research areas. Statistical variable selection methods cannot distinguish between confounders, mediators, and colliders without causal assumptions, potentially leading to biased estimates. Manual literature review becomes increasingly impractical as PubMed now contains over 35 million citations and adds approximately 1.5 million new articles annually.

Recent advances in causal inference methodology have emphasized the importance of directed acyclic graphs (DAGs) for representing causal assumptions and identifying appropriate adjustment sets.^12,16^ DAG-based approaches offer a principled framework for distinguishing between different types of bias and determining the minimal sufficient sets of variables required for confounding control^18^. However, constructing realistic DAGs for complex biomedical phenomena remains challenging when attempting to incorporate the breadth of relevant causal knowledge documented in the scientific literature. This challenge is compounded by four fundamental barriers: overwhelming literature volume, inconsistent terminology across publications, ambiguous directionality of reported relationships, and lack of systematic approaches for integrating literature-derived knowledge into formal causal models.

To address these barriers, we require computational approaches that can automatically extract, validate, and synthesize causal knowledge from large-scale literature repositories. The Semantic MEDLINE Database (SemMedDB) provides a foundation for such approaches, containing over 100 million semantic predications extracted from PubMed abstracts using the SemRep natural language processing system. ^7^ These structured subject-predicate-object triples capture causal, associational, and mechanistic relationships between biomedical concepts. Unlike manually curated databases with limited coverage, SemMedDB offers comprehensive scope across biomedical domains while maintaining a structured format suitable for automated processing.

However, raw extraction from SemMedDB presents challenges including low-quality predications, contradictory relationships, and terminological inconsistencies requiring systematic filtering and validation. Recent developments in large language models offer promising solutions for enhancing the quality and interpretation of literature-derived causal knowledge. ^19^ Building on these capabilities, we developed CausalKnowledgeTrace to systematically address each barrier to literature-based variable selection through automated extraction, multi-stage quality filtering, semantic consolidation, large language model-based validation, and systematic bias detection.^2^

The primary objectives of this work are twofold. First, we develop a scalable computational framework for constructing literature-derived directed acyclic graphs that systematically inform variable selection in causal studies while maintaining complete transparency about evidentiary basis. Second, we create user-friendly interfaces and standardized output formats that facilitate seamless integration of literature-derived causal knowledge into existing epidemiological workflows.

The remainder of this paper describes the technical implementation of CausalKnowledgeTrace, evaluates its performance through comprehensive comparison with expert judgment and systematic literature review, and discusses the implications for evidence-based causal inference practice in biomedical research.

## 2. Methods

### 2.1 System Overview

CausalKnowledgeTrace is a modular computational framework integrating Python-based analysis components with a Django web interface. The system utilizes the Semantic MEDLINE Database (SemMedDB) as its primary knowledge source, containing structured semantic predications extracted from PubMed abstracts. Graph construction leverages pre-generated causal assertion files in JSON format with directed relationships between biomedical concepts, each including supporting PubMed identifiers (PMIDs). The analysis pipeline is implemented using NetworkX for graph operations, following six systematic stages that transform literature-derived causal assertions into validated directed acyclic graphs suitable for formal causal inference.

**Figure 1.**
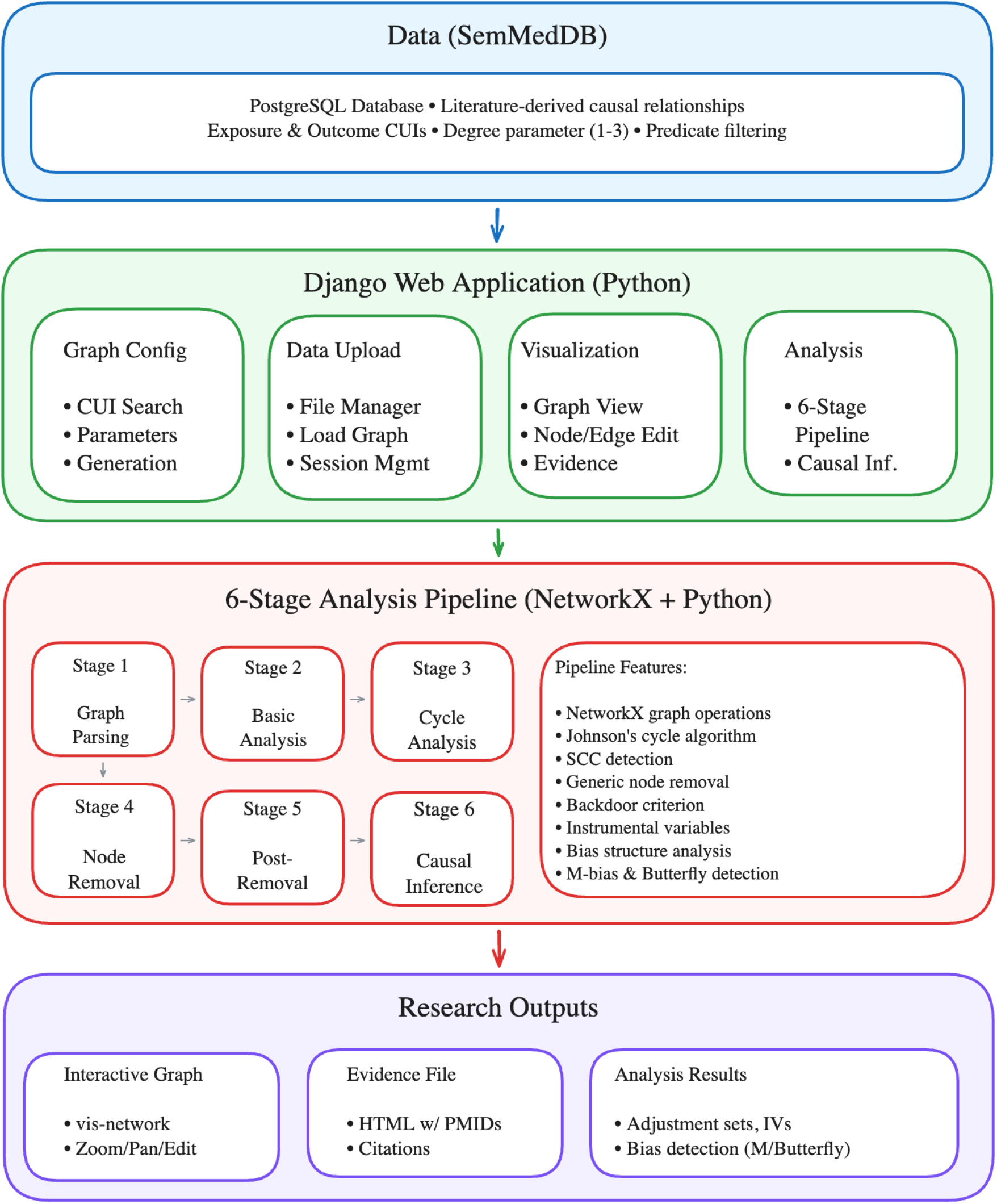
CausalKnowledgeTrace workflow and system architecture.

### 2.2. Analysis Pipeline

Each stage is implemented as an independent Python module with standardized input/output interfaces.

#### 2.2.1. Graph Construction

The pipeline converts JSON-formatted causal assertions into NetworkX directed graph structures, preserving complete metadata including semantic predicates (e.g., CAUSES, AFFECTS), supporting PubMed identifiers (PMIDs).

#### 2.2.2. Graph Characterization

The pipeline computes node-level metrics (in-degree, out-degree, betweenness centrality, closeness centrality) and graph-level statistics (density, diameter, strongly connected components). Betweenness centrality identifies structural bottlenecks representing critical mediators or artifactual hub nodes, informing subsequent quality control decisions.

#### 2.2.3. Cycle Detection

Directed acyclic graph (DAG) compliance is essential for formal causal inference. It implements exhaustive cycle detection using Johnson’s algorithm and identifies strongly connected components using Tarjan’s algorithm. The pipeline tracks per-node cycle participation frequencies, cycle length distributions, and representative cycles for detailed inspection.

#### 2.2.4. Graph Refinement

Literature-derived causal graphs frequently contain high-centrality generic terms (e.g., “Disease”, “Symptoms”) that introduce artifactual relationships and spurious cycles. It implements systematic removal of such nodes through combined centrality analysis and expert curation. The pipeline maintains a curated list of generic biomedical terms and computes the cycle reduction achieved by their removal. Each node is removed along with its incident edges, and the resulting graph is re-analyzed for cycle content, preserving clinically meaningful causal relationships while eliminating artifactual structures.

#### 2.2.5. Post-Refinement Validation

Following the removal of generic nodes, it recomputes metrics from Stages 2 and 3 to quantify residual cycles and assess progress toward DAG compliance. Nodes are ranked by residual cycle participation frequency. If DAG requirements are not satisfied, the pipeline generates recommendations for further intervention, prioritizing nodes with high cycle participation but lower clinical relevance.

#### 2.2.6. Causal Inference and Bias Detection

The final stage applies formal causal inference methods to DAG-compliant graphs for user-specified exposure and outcome concepts.

##### Confounder Identification

A node *C* qualifies as a confounder candidate if (*C → A*) *∈ E* and (*C → Y*) *∈ E* for exposure *A* and outcome *Y* in graph *G* = (*V, E*). The pipeline applies the backdoor criterion to identify minimally sufficient adjustment sets.

##### Bias Structure Detection

The system identifies (1) *M-bias* structures involving common causes of variables related to exposure or outcomes; (2) *Butterfly bias* where confounders also function as colliders along backdoor paths; and (3) *Colliders* representing common effects that should be excluded from adjustment sets.

##### M-bias Detection

The pipeline detects M-bias structures systematically by evaluating direct (1-hop) parent-child relationships, enabling robust detection in the graph produced by the CausalKnowledgeTrace application. A node *M* is classified as an M-bias collider if it is a common direct child of different parents of exposure *A* and outcome *Y*. A node is called an M-bias collider if the following holds: given an exposure *A*, an outcome *Y*, and two distinct nodes *P*_1_ and *P*_2_ such that (*P*_1_ *→ A*) *∈ E* and (*P*_2_ *→ Y*) *∈ E*, a node *M* is a collider if it satisfies (*P*_1_ *→ M*) *∈ E* and (*P*_2_ *→ M*) *∈ E*, provided that *M ∈/* {*A, Y*} *∪ C* (where *C* is the set of previously identified confounders). The algorithm strictly ensures that *M* is neither the exposure nor the outcome, nor a previously identified confounder. By isolating these specific 5-node structures *A ← P*_1_ *→ M ← P*_2_ *→ Y*, the system automatically flags these colliders, allowing researchers to avoid them in adjustment sets and thereby prevent the unintended opening of spurious backdoor paths.

##### Butterfly Bias Detection

Butterfly bias arises when a valid confounder is also a collider of two or more independent confounders, thereby creating a spurious backdoor effect. To identify these topological configurations, the pipeline examines the local neighborhoods of all previously identified confounders. For each valid confounder *C*, the algorithm identifies the direct predecessor for each valid confounder *C* and intersects these sets with the global pool of known confounders. The node *C* is designated as a butterfly confounder if it is the direct child of at least two distinct confounders (for example, *C*_1_ and *C*_2_). The system automatically flags hazardous variables, allowing researchers to avoid directly adjusting the butterfly node and instead adjust for its parent confounders to mitigate collider stratification bias.

#### Confounder Classification

Confounders are classified as follows: (1) *Independent* (no confounder parents); (2) *Single-parent* (one confounder parent); (3) *Butterfly* (multiple confounder parents requiring specialized adjustment strategies).

### 2.3. Implementation and Performance

The pipeline maintains complete provenance tracking, preserving metadata, including semantic predicates, and supports PubMed identifiers (PMIDs). Each edge is traceable to its supporting literature, enabling validation that graph-theoretic confounders correspond to substantive causal claims.

The system provides a Django-based web application that integrates the Python pipeline with interactive visualization using PyVis. Key features include: (1) interactive graph upload and validation, (2) real-time pipeline execution with progress monitoring, (3) interactive visualization with node/edge filtering, (4) comprehensive results dashboard with downloadable outputs, and (5) literature provenance exploration with direct PubMed linking.

The pipeline demonstrates excellent computational efficiency. For the hypertension-Alzheimer’s analysis, processing times ranged from 0.3 to 1.0 seconds across all three degree levels (degree 1: 361 nodes, 429 edges; degree 2: 727 nodes, 1,215 edges; degree 3: 866 nodes, 1,442 edges). Memory usage scales linearly with graph size.

### 2.4. Evaluation Framework

To validate CausalKnowledgeTrace, we analyzed the well-studied relationship between hypertension and Alzheimer’s disease across three degree neighborhoods (1-3). The system constructed causal graphs at 1-degree (361 variables, 429 relationships), 2-degree (727 variables, 1,215 relationships), and 3-degree (866 variables, 1,442 relationships), examining graph complexity scaling, cycle detection and resolution, generic node removal effectiveness, and confounder identification.

## 3. Results

CausalKnowledgeTrace analysis of the hypertension–Alzheimer’s disease relationship demonstrated systematic scaling across degree neighborhoods: 361 variables/429 relationships (degree 1), 727 variables/1,215 relationships (degree 2), and 866 variables/1,442 relationships (degree 3). Graph density decreased with expansion (0.0033, 0.0023, 0.0019), indicating increasingly sparse networks.

The analysis revealed complex cyclic structures with baseline cycle counts of 54, 602, and 606 across degree levels, with 37-56 nodes (6.5-10.2%) participating in cyclic relationships. Key confounders identified across all degrees included inflammation, diabetes, insulin resistance, obesity, and ischemia. Processing times ranged from 0.3 to 1.0 seconds, demonstrating computational efficiency.

**Table 1.** Hypertension–Alzheimer’s Causal Graph Analysis Results by Degree Level

| Metric | Degree 1 | Degree 2 | Degree 3 |
| --- | --- | --- | --- |
| Total Variables | 361 | 727 | 866 |
| Causal Relationships | 429 | 1,215 | 1,442 |
| Graph Density | 0.0033 | 0.0023 | 0.0019 |
| Nodes in Cycles | 37 (10.2%) | 52 (7.2%) | 56 (6.5%) |
| Baseline Cycles<sup>b</sup> | 54 | 602 | 606 |
| Processing Time | 0.3s | 0.7s | 1.0s |

<sup>a</sup>Number of strongly connected components containing cycles
<sup>b</sup>Total count of simple cycles in the graph

### 3.1. Confounding Bias

Structural analysis of the CausalKnowledgeTrace-derived directed graph identified several key upstream common causes, highlighting them as significant sources of confounding bias. Specifically, the algorithm identified major cellular, metabolic, and lifestyle factors—namely **obesity, oxidative stress, injury, dietary intervention**, and **physical activity**—as structurally acting as confounders in the relationship between Hypertension and Alzheimer’s disease. The algorithmic identification of these specific pathways strongly aligns with established epidemiological and pathophysiological consensus. The literature widely recognizes these variables as known, critical confounders that independently exacerbate, or, in the case of lifestyle interventions, structurally modulate both cardiovascular risk and neurodegenerative decline^1,9,11,13^.

**Figure 2.**
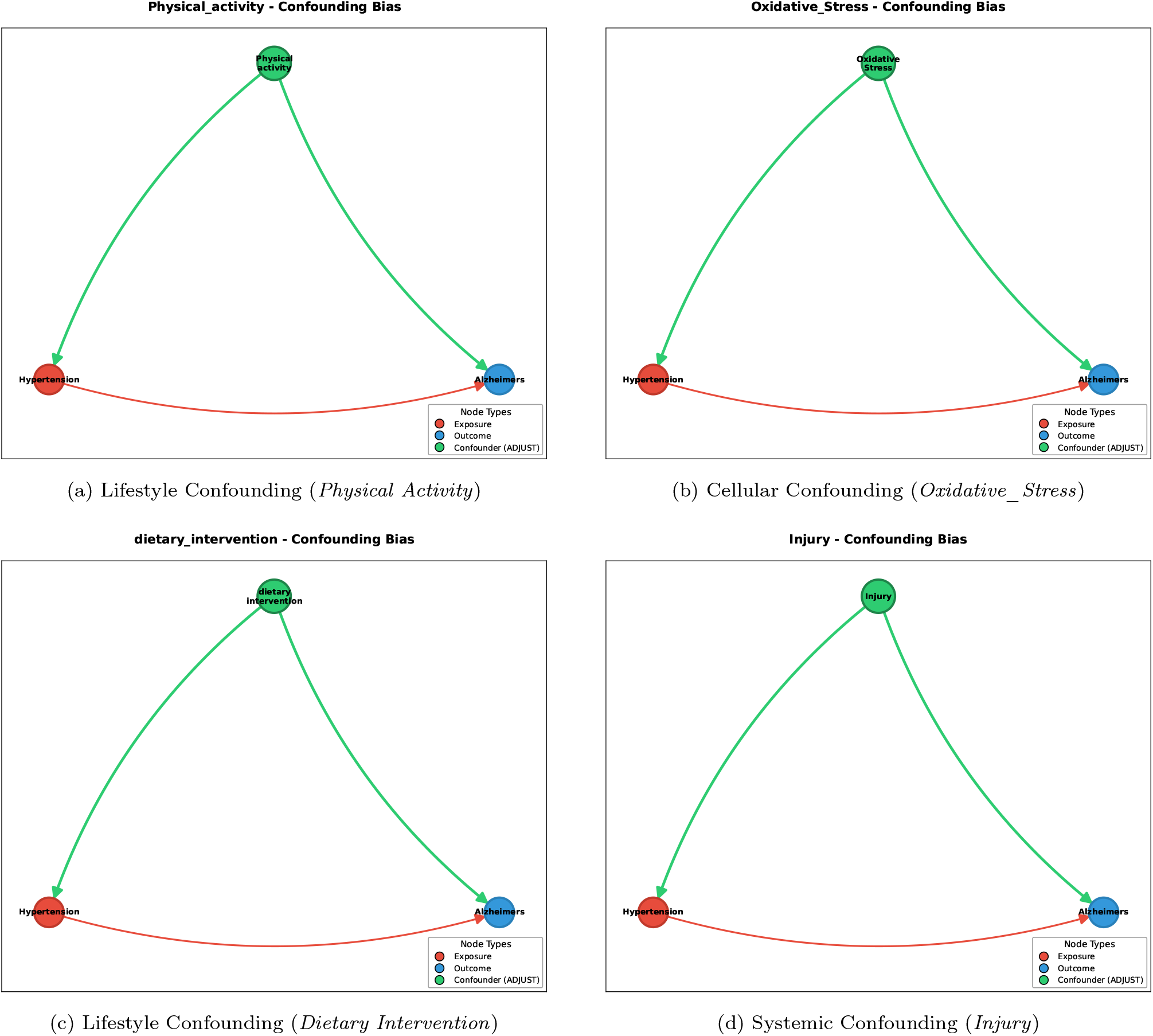
Algorithmic isolation of Confounding Bias (*E ← X → O*). The CausalKnowledgeTrace pipeline successfully extracted upstream common causes across all major physiological domains. Variables such as metabolic disease parameters (Obesity), exogenous lifestyle modifications (Dietary intervention and Physical Activity), acute physiological stress (Injury), and intracellular damage (Oxidative stress) were consistently identified as structural confounders that strictly necessitate statistical adjustment.

### 3.2. Mediators

Subsequent analysis of the CausalKnowledgeTrace-derived graph identified direct patho-physiological pathways linking hypertension to Alzheimer’s disease. The algorithm successfully flagged eleven variables as structural mediators on the causal path (Hypertension *→* Mediator *→* Alzheimer’s). Notably, it extracted nodes signifying cerebrovascular and cellular damage mechanisms: specifically *Ischemia, Hypoxia, Reactive Oxygen_Species*, Nitric Oxide, *Inflammation*, and *Vascular Diseases*. By graphically situating these nodes along the causal path, our structural model correctly reflects current neurovascular literature. Clinical research extensively demonstrates that chronic hypertension physically alters cerebral vasculature, inducing a state of chronic hypoperfusion (*Ischemia* and *Hypoxia*) and oxidative damage (*Reactive Oxygen Species*), which directly accelerates the accumulation of amyloid-beta^6,22^. If researchers adjust for these intermediate effects in statistical modeling, they will effectively block the causal pathway, leading to overadjustment bias that artificially masks the true total effect of hypertension on dementia risk^14^.

**Figure 3.**
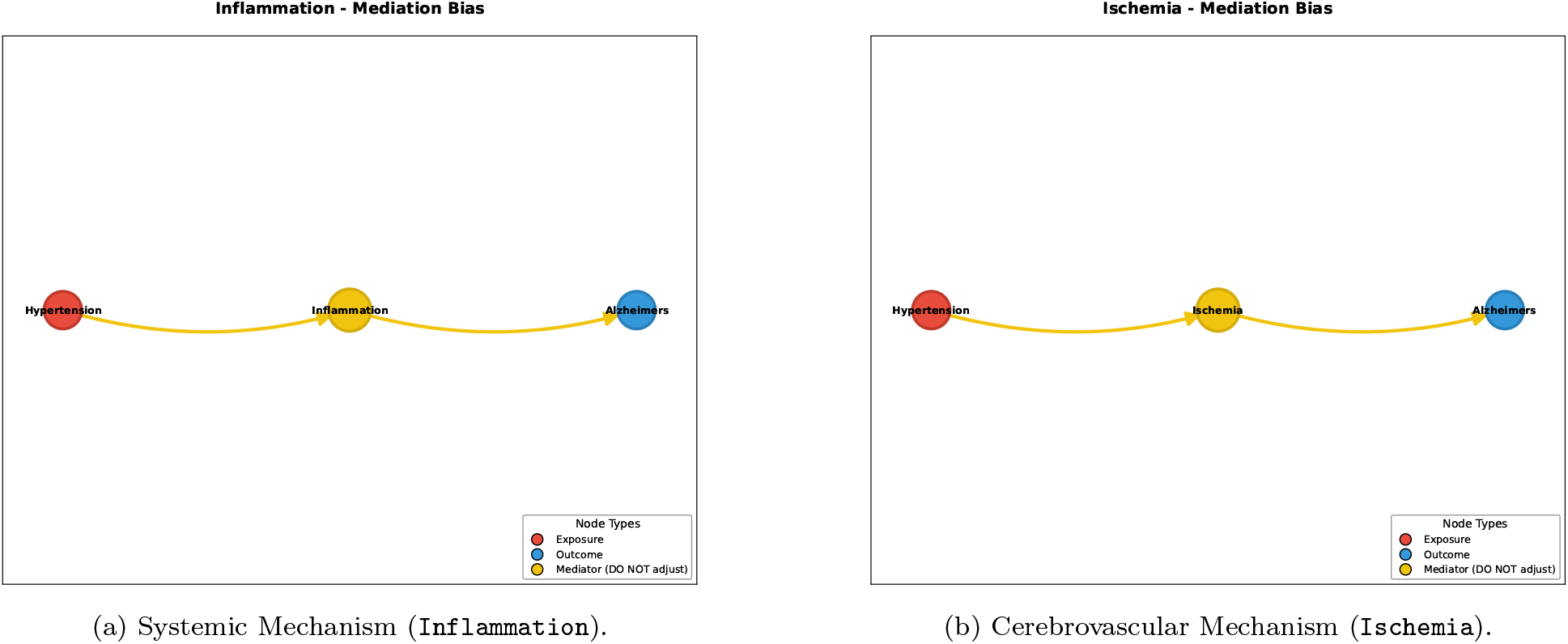
Structural extraction of Mediation Bias (*E → X → O*). The CausalKnowledgeTrace pipeline identified core pathophysiological pathways bridging hypertension and Alzheimer’s disease. Systemic mediators (Inflammation) and direct cerebrovascular damage markers (Ischemia) lie strictly on the causal path between the exposure and the outcome. Statistically adjusting for these intermediate biological endpoints would induce severe overadjustment bias, artificially blocking the true total effect of vascular disease on neurodegeneration.

### 3.3. Colliders

CausalKnowledgeTrace structural analysis identified downstream consequences of both hypertension and Alzheimer’s disease neuropathology, flagging variables that pose a risk of collider-stratification bias if conditioned on during analysis. Algorithmically classified colliders included widespread neuronal damage (`Neurons`), generalized disability (`Disability_NOS`), and mortality (`Cessation_of_life`). Each variable represents a clinical endpoint with independent causal pathways from severe hypertensive cardiometabolic disease and advanced Alzheimer’s neuropathology, forming structures of the type Hypertension *→* Mortality *←* Alzheimer’s disease. Conditioning on such variables (for example, by restricting the analysis to individuals who survived to advanced age or who exhibited end-stage functional decline) opens a non-causal path between otherwise independent exposures, inducing spurious associations. This phenomenon, collider stratification bias, is well-documented in dementia epidemiology, where conditioning on survival or shared downstream functional impairment artificially inverts associations between established risk factors. CausalKnowledgeTrace identification of these structures enables investigators to avoid analytic conditioning on shared consequences prior to study design^3,10,21^.

### 4. Discussion

CausalKnowledgeTrace automates the construction of causal graphs from biomedical literature, reducing reliance on expert-driven variable selection and enabling principled, transparent confounder identification and causal modeling at scale. The tool draws on relationships extracted from millions of published abstracts, translating accumulated epidemiological evidence into executable graph structures that support rigorous observational study design.

### 4.1. Key Findings and Technical Contributions

Our evaluation demonstrated substantial agreement between the causal graphs generated by CausalKnowledgeTrace and expert knowledge while identifying additional relationships overlooked in systematic reviews and detecting sophisticated bias patterns. Existing computational approaches to causal discovery typically fall into two categories: constraint-based algorithms that learn causal structure from data and knowledge-based methods that rely on expert-curated databases^12^. Constraint-based methods often struggle with the sample-size requirements and assumptions needed for reliable causal discovery in biomedical settings. Expert-curated approaches offer high-quality knowledge but are limited by their restricted coverage and infrequent updates. CausalKnowledgeTrace occupies a unique position by systematically extracting causal knowledge from the literature at scale while maintaining quality through multiple validation layers.

The semantic consolidation module automatically identifies and merges synonymous concepts while preserving complete provenance information. The integration of large language models for relationship validation provides contextual assessment of directionality and appropriateness. The modular architecture supports both standalone analysis and integration into larger computational workflows. Traditional approaches to variable selection in causal inference rely heavily on domain expertise, statistical screening methods, or simple directed acyclic graphs constructed through expert consensus^15,20^. These methods may overlook documented confounding relationships, lack systematic approaches to bias identification, and provide limited traceability to supporting evidence. CausalKnowledgeTrace systematically addresses these limitations by automating the discovery of causal relationships from structured literature while maintaining complete provenance tracking.

The identification of complex bias patterns represents a significant advancement^2^. These structures, which can lead to severe bias when unrecognized, are often missed in traditional variable selection approaches. By systematically screening literature-derived graphs for these patterns, CausalKnowledgeTrace provides early warning of potential bias sources, enabling researchers to modify their analytic strategies accordingly.

### 4.2. Limitations and Future Directions

CausalKnowledgeTrace’s reliance on published literature means that emerging risk factors or relationships documented primarily in gray literature may be underrepresented. Publication bias^17^ may skew the identification of protective factors compared to risk factors. The system’s focus on semantic predications extracted from abstracts^7^ may overlook important contextual information that appears only in full-text articles. Additionally, relationships well documented in the literature but based on methodologically flawed studies may receive high confidence scores, underscoring the importance of combining automated knowledge extraction with critical appraisal of study quality. Because SemMedDB aggregates predications uniformly across all available literature without weighting contextual directionality or study quality, contradictory findings and pure negative observations contribute equally to the knowledge graph. This automated aggregation frequently interprets explicit negative experimental findings as positive causal edges—such as extracting a *Glucose → Insulin* edge from a study documenting that insulin release remained unaffected ^8^—generating incredibly dense, spurious bidirectional edges between biomedical concepts that introduce inescapable and meaningless feedback cycles.

To address these limitations and enhance CausalKnowledgeTrace’s capabilities, several avenues for future development are planned. A key priority is the development of an LLM-based system for directly extracting causal relationships from biomedical literature. This system would automatically process newly published articles, extract relevant causal predications, validate their quality and relevance, and update the underlying knowledge base in real-time. The LLM-based extraction pipeline would complement existing SemMedDB predications by providing more comprehensive coverage and reducing reliance on periodic database releases.

Additional future directions include integrating with biological pathway databases and clinical trial registries to provide more comprehensive causal knowledge, developing population-specific filtering capabilities to construct causal graphs tailored to specific demographic groups or clinical populations, and enhancing human-in-the-loop capabilities to support domain experts. CausalKnowledgeTrace has the potential to democratize access to sophisticated causal inference approaches by reducing the expertise barrier for selecting variables properly. The emphasis on transparency and provenance tracking aligns with broader trends toward reproducible and interpretable research, supporting the verification and validation of analytical decisions.

## 5. Conclusion

CausalKnowledgeTrace represents a novel Python-based computational framework that systematically extracts and analyzes causal knowledge from biomedical literature to inform variable selection in complex observational studies. Our analysis of the hypertension-Alzheimer’s relationship demonstrates the system’s ability to systematically scale across causal neighborhoods, identifying 361-866 variables with 429-1,442 causal relationships, along with comprehensive cycle analysis and bias detection.

The six-stage Python pipeline provides comprehensive cycle analysis (54-606 cycles across degree levels) and systematic generic node removal capabilities, enabling the identification of clinically meaningful confounders, including inflammation, diabetes, insulin resistance, obesity, and ischemia. Processing efficiency (0.3-1.0 seconds for all three degrees) demonstrates computational scalability for large-scale biomedical applications.

Key technical innovations include comprehensive provenance tracking, automated bias structure detection, and integration of formal causal inference methods with literature-derived evidence. The Django web interface provides user-friendly access while maintaining the computational rigor of the underlying Python pipeline.

Essential limitations include reliance on published literature and abstract-level semantic predications, emphasizing that the system should complement rather than replace expert judgment. However, the systematic approach to causal graph construction provides a principled foundation for evidence-based variable selection that scales with the expanding biomedical knowledge base.

## Acknowledgments

This work was supported by the National Library of Medicine under Award Number R00LM013367. The content is solely the responsibility of the authors and does not necessarily represent the official views of the National Institutes of Health. We thank the University of New Mexico Department of Computer Science and the Health Sciences Center for computational resources and technical support. Additionally, we are grateful to the National Library of Medicine for providing access to SemMedDB and UMLS resources.

## Data Availability

CausalKnowledgeTrace is freely accessible as a web application at https://habanero.health.unm.edu/CausalKnowledgeTrace/. The source code is available at https://github.com/unmtransinfo/CausalKnowledgeTrace. The SemMedDB database is publicly available at https://skr3.nlm.nih.gov/SemMedDB/. Analysis scripts and supplementary materials are available upon request. A UMLS license is required for database access and can be obtained free of charge from the National Library of Medicine.

